# Combining RNA-seq and proteomics boosts sensitivity to detect differentially expressed genes by rpMeta

**DOI:** 10.64898/2026.10.15.758995

**Authors:** Xijun Zhang, Joseph P. LaMorte, Robert L. Kortum, Craig D. Shriver, Clifton L. Dalgard, Matthew D. Wilkerson

## Abstract

Differential gene expression analysis is a powerful technique to characterize biological systems. Differentially expressed genes (DEGs) are commonly identified by analysis of RNA sequencing datasets or mass spectrometry proteomics datasets. Increasingly, multi-omic studies include both RNA sequencing and proteomics from the same biospecimens. However, the common multi-omic DEG analysis practice involves separate DEG analysis by RNA and by protein with limited comparison by fold change correlations or Venn Diagrams. This disjoint practice may not utilize all of the available signal from both datatypes as transcripts and proteins have correlated expression levels. To integrate RNA and proteomics, we developed a new method, RNA protein meta-analysis called rpMeta. rpMeta takes RNA and protein expression data and calculates DEGs by RNA, by protein, and by meta-analysis models, following user-specified options. Spike-in, downsampling and cross-cohort benchmarks revealed that the meta-analysis model had greater sensitivity than RNA and protein models. In a real-world use case, the meta-analysis model discovered new genes and biological processes in lung cancer subtypes that were not detectable by the RNA or protein models. To streamline interpretation, rpMeta produces publication-ready data visualizations of the integrated DEGs. rpMeta is an open-source R package, available at https://github.com/zhangx8/rpMeta.

**GRAPHICAL ABSTRACT:** 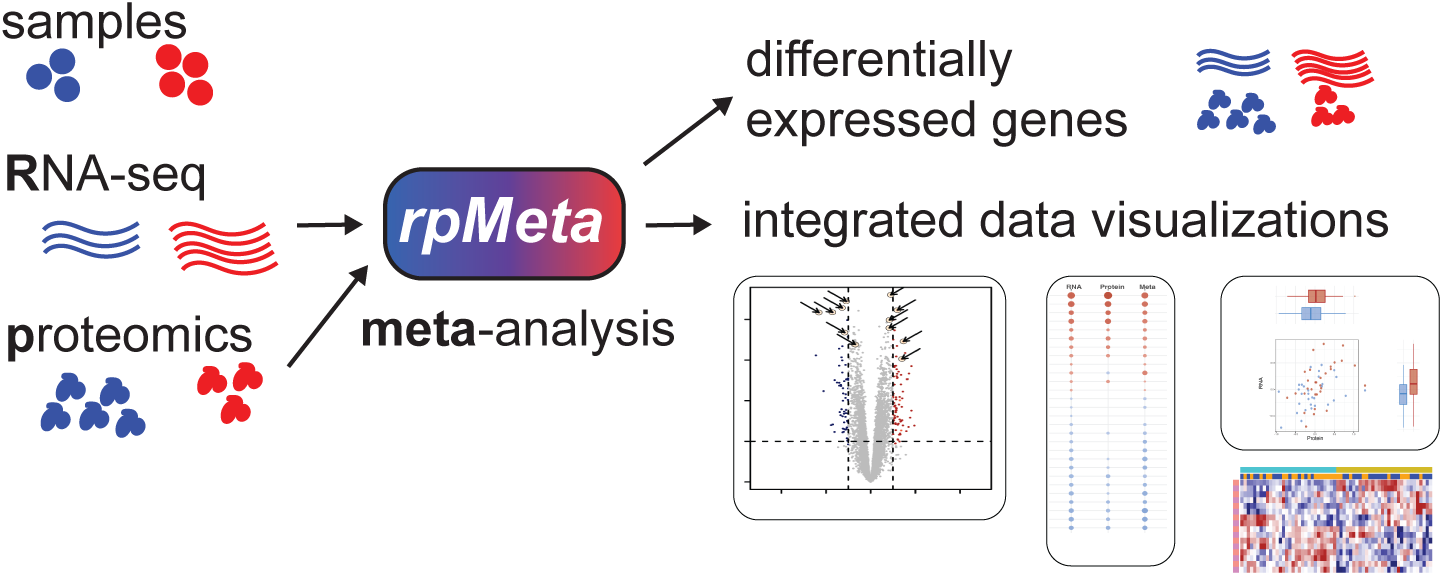

## INTRODUCTION

Genome-wide gene expression analysis has been a revolutionary method in biological research, resulting in the discovery of differentially expressed genes (DEGs), across a myriad of biological systems (1,2). These DEGs have fueled many biomedically-relevant discoveries such as expression signatures of response to cancer immunotherapy (3) and response to antidepressants (4). The predominant laboratory techniques for genome-wide gene expression analysis are the sequencing of RNA transcripts (RNA-seq) and the bottom-up mass spectrometry quantification of proteins (MS-based proteomics) (5,6). Many methods for DEG analysis have been developed for both platforms. However, leading tools and workflow options such as gene filtering and normalization are known to affect DEG results and can underestimate DEGs in the data that may be statistically and biologically significant (2,7).

Recently, multi-omic studies combine multiple analytic platforms, such as RNA-seq and MS-based proteomics, to gain broader views of biological systems. Large cohort multi-omic studies include The Genotype-Tissue Expression Project (GTEx) (8,9), The Cancer Genome Atlas (TCGA) (10), the Clinical Proteomic Tumor Analysis Consortium (CPTAC) (11), and the Applied Proteogenomics Organizational Learning and Outcomes network (APOLLO) (12–15). To date, the common DEG analysis approach in multi-omic studies involves separate DEG analysis by RNA-seq and by proteomics and then comparing results such as by fold change correlations, Venn diagrams or descriptive text (16–21). However, this disjoint approach does not maximize the potential joint signal available from both RNA and protein. RNA transcription and protein translation are sequential steps in gene expression process, for protein-coding-genes. As many multi-omics studies have revealed that the vast majority of RNAs and proteins have positively correlated gene expression levels (8,13,22), there is potential additional signal of differential expression from the combined RNA and protein that is greater than the signal by RNA alone or protein alone. This additional signal could power the detection of DEGs with subtle changes in expression and in studies with small sample sizes.

A powerful approach to combining expression datasets is meta-analysis (23), which can increase statistical power. Meta-analysis methods include different ways of combining p-values or effect sizes across datasets to calculate a meta-analysis p-value (24). Meta-analysis methods have been widely used with RNA datasets from RNA sequencing or microarrays (25,26) and also with MS-based proteomics datasets (27,28). At this time, meta-analysis DEG approaches have not been applied or evaluated in combining RNA-seq and proteomics datasets together, so the performance of DEG meta-analysis of RNA-seq and proteomics data is unknown. No specialized analysis tool for combining RNA-seq and proteomics in DEG meta-analysis is currently available. Additionally, data visualization in multi-omics can be complex and is critical to identify and comprehend patterns in the data (5,29). To date, there is not a specialized visualization tool available for multi-omic DEG analysis.

Here, we describe our novel method and tool for RNA and protein meta-analysis, called rpMeta. rpMeta is an open-source R package under the MIT license. rpMeta combines genome-wide RNA-seq and proteomics datasets and detects differentially expressed genes by meta-analysis statistics. rpMeta is highly customizable for the analyst, such as through differential expression methods, meta-analysis methods, and other options. rpMeta includes custom data visualizations for multi-omic DEG analysis and generates publication-quality figures. To assess if rpMeta increases performance of gene expression analysis compared to RNA-only or protein-only analysis, we performed three benchmarks, a spike-in simulation, a downsampling analysis, and a cross-cohort analysis. For these benchmarks, we compared two well-known subtypes of lung adenocarcinoma, Terminal Respiratory Unit (TRU) and Proximal-Proliferative (PP) subtypes (17,30–34), in published RNA-seq and proteomics datasets from the CPTAC (17) and APOLLO cohorts (13). These subtypes have distinct biology in lung cancer (13,30,35,36) and are known to have large numbers of differentially expressed genes. These benchmarks demonstrated the superior performance of the meta-analysis (meta) model, compared to the RNA or protein models. Finally as a real-world use case, we applied rpMeta to all TRU and PP samples from the APOLLO cohort and discovered 376 genes as well as novel biological processes that were not detected by RNA or protein models and were relevant to their corresponding subtype’s biology.

## MATERIALS AND METHODS

### Meta-analysis of RNA-seq and MS-based proteomics by rpMeta

rpMeta is a bioinformatics analysis tool for identifying differentially expressed genes from RNA-seq and proteomics datasets from the same matched specimens. rpMeta is written in R, is open source, and includes a detailed tutorial and documentation. rpMeta is also the main driver function for the package and has 3 required input files (Figure 1). First, the RNA expression matrix is gene by sample matrix of RNA transcript expression derived from RNA sequencing. The RNA expression values can be either read counts, transcripts per million, or other normalized values such as upper quartile normalized counts. Second, the protein expression matrix is a gene by sample matrix of protein expression derived from MS-based proteomics. The protein values are normalized ratios or values of protein expression. Both the RNA and protein expression data are required to correspond to the same gene database, for example HUGO gene symbols, which serve as row names. Third, a sample information file defines sample groups and covariates for each sample. Additional user options define which groups to compare and possible covariates.

**Figure 1:**
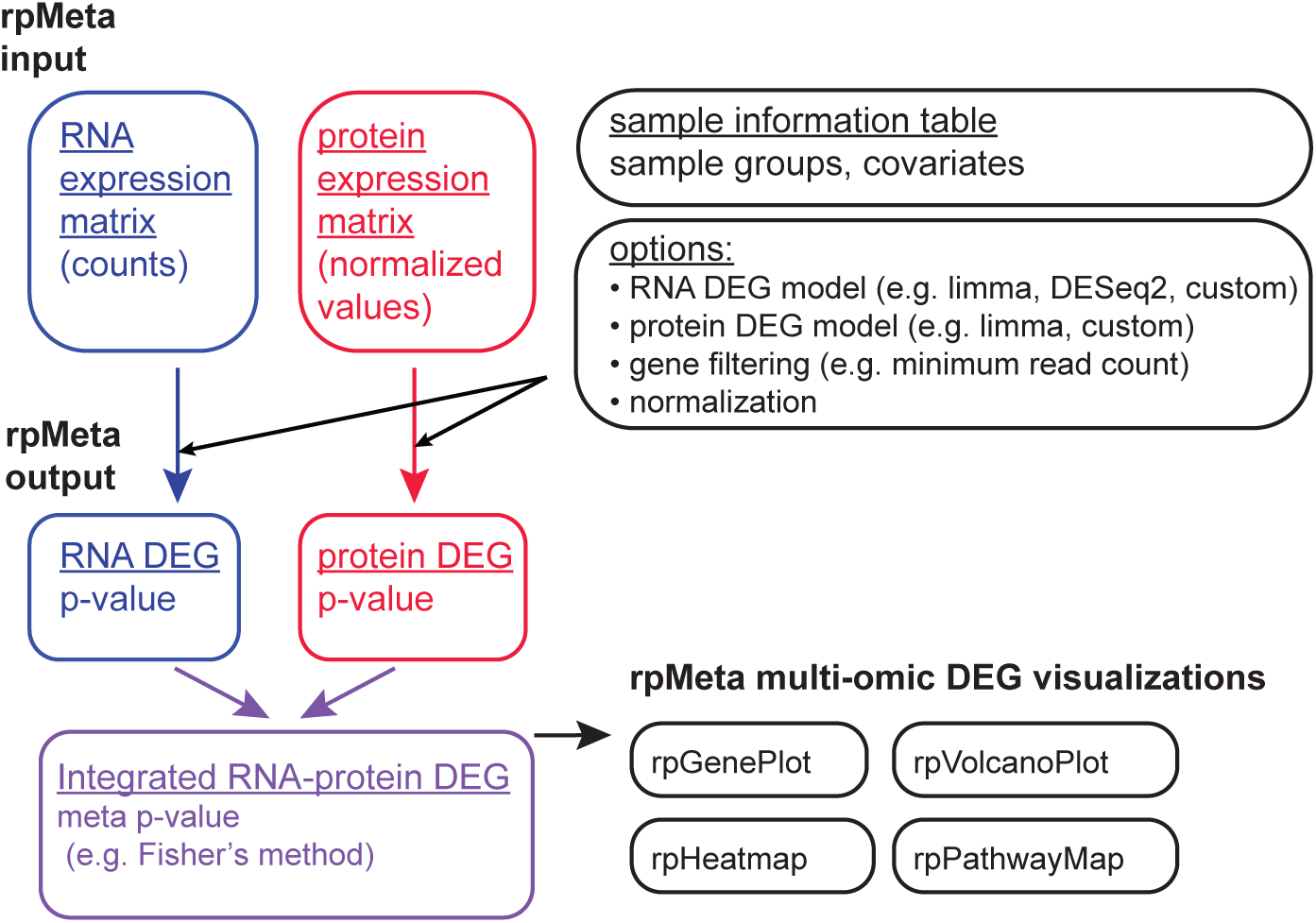
Design of rpMeta method. This figure displays the input, methods, output and visualization utilities of rpMeta.

With those input data, rpMeta filters genes by user options for minimum expression and maximum proportion of missing values. rpMeta has two modes for DEG analysis, a two group comparison mode and a mode for expression association with a continuous variable. For the two group mode, rpMeta calculates DEGs in the RNA expression data by calling either DESeq2 (37), limma (38) or a custom DEG R function as provided by the user. The custom DEG function provides flexibility for users to easily extend rpMeta with new differential expression statistics, such as a Wilcoxon Rank Sum Test example DEG function included in the rpMeta tutorial. This custom DEG function is conceptually similar to custom R function that we provided in another package, ConsensusClusterPlus, which has been highly adopted by the bioinformatics research community (39). rpMeta also includes an option to normalize RNA prior to calling limma, such as voom (38). Then, rpMeta calculates DEGs in the protein expression data by calling limma or a custom DEG function, which could be the same or a different function than the RNA DEG method. For the continuous variable mode, rpMeta tests for Spearman correlation of RNA and protein expression with a user-supplied continuous variable.

Next, rpMeta combines p-values between RNA DEGs and protein DEGs by a meta-analysis method. rpMeta has the default meta-analysis method of Fisher’s method of combining p-values, as well as additional methods of Pearson’s, Stouffer’s, a Weighted Stouffer’s Method, Tippett’s method, and Wilkinson’s method (Figure 1) (24). For genes only detected by RNA-seq, the meta p-value is set to the RNA p-value and similarly for genes only detected by proteomics. False discovery rates for each model, RNA, protein, and meta, are calculated by the Benjamini-Hochberg (40) method by p.adjust. Finally, rpMeta combines the DEG statistics for RNA, protein and meta models into publication ready integrated DEG tables which are written to text files, one table for common genes (Table S1A,B) and another table for all genes from both platforms.

### Multi-omic expression visualization in rpMeta

To enable analyst and reader interpretation of multi-omic differential expression, rpMeta provides novel data visualization utilities. Two utilities provide a visualization of the global properties of DEGs across RNA, protein and meta models. For a brief summary, the rpVenn utility plots a multi-way Venn diagram of DEGs across the three models (Figure 2A). The rpVolcano utility displays two Volcano plots (41) for RNA and protein models separately, and also displays RNA and protein fold changes in a fold change scatter plot (Figure 2B). rpVolcano enables a user to comprehend the signal alignment of differential expression between RNA and protein and to see individual genes in which the signal may be stronger on the RNA side or protein side.

**Figure 2:**
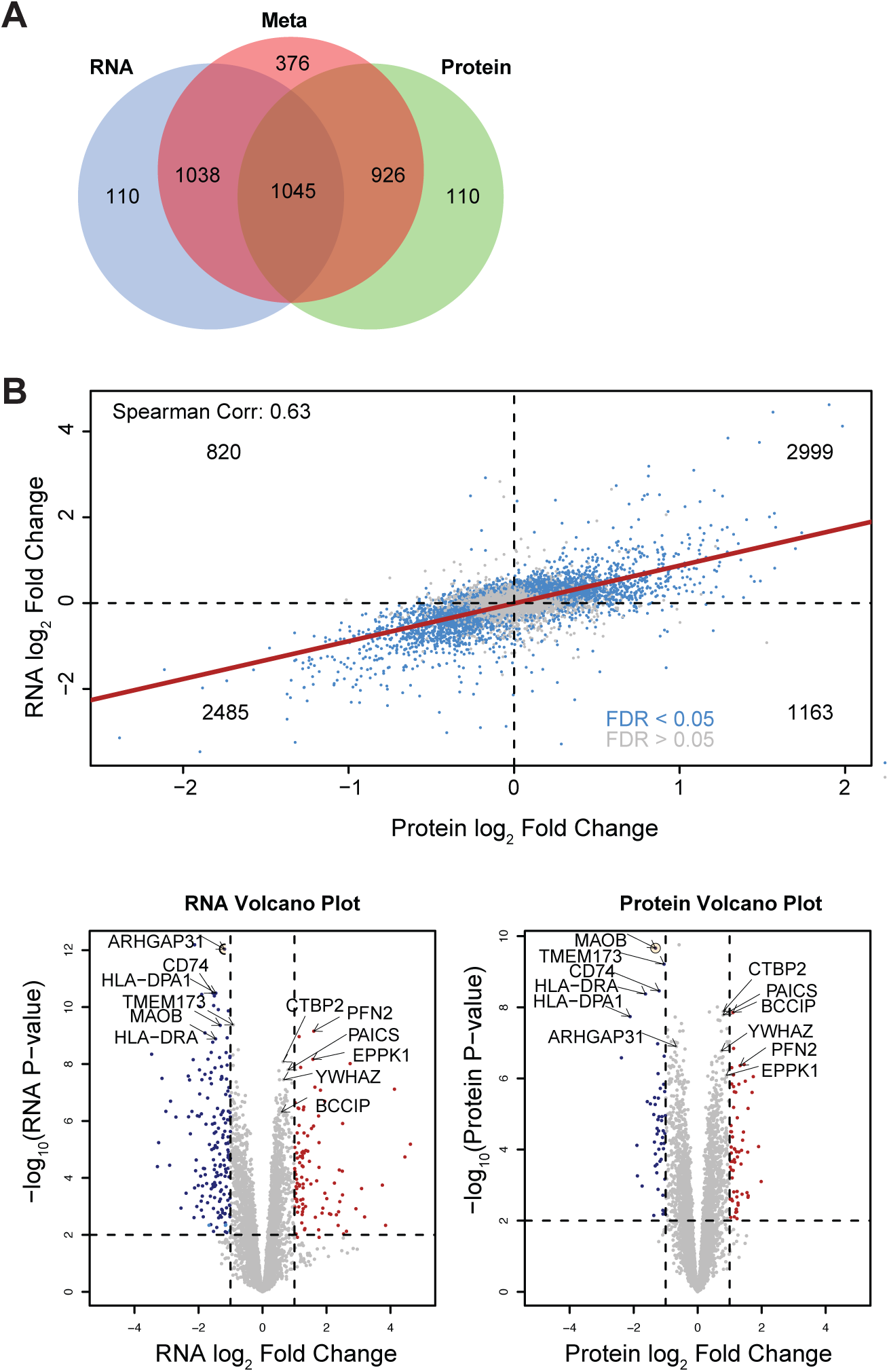
Data visualizations of rpVolcano, joint fold change, and rpVenn. Visualizations for differential expression between TRU and PP subtypes, n = 30 and n =30, respectively, from the APOLLO cohort. Numbers of DEGs and overlaps between models of rpMeta displayed as a Venn Diagram (A). The scatter plot displays log2 fold changes between protein and RNA (B). rpVolcano plots display fold change versus p-values for RNA and protein (B).

To enable focused analysis on individual genes, the rpGenePlot utility provides a scatter plot of RNA and protein expression and combined with RNA-specific and protein-specific boxplots for genes of interest (42) (Figure 3A,B). For a given gene, the rpGenePlot provides a clear visualization of the expression relationship between RNA and protein and the differential expression between sample groups by RNA and by protein. In this way, rpGenePlot provides effective, publication-ready illustrations of differential expression for key genes of interest for multi-omic analysis publications. rpMeta also includes similar utilities for the continuous DEG mode as described in the rpMeta tutorial.

**Figure 3:**
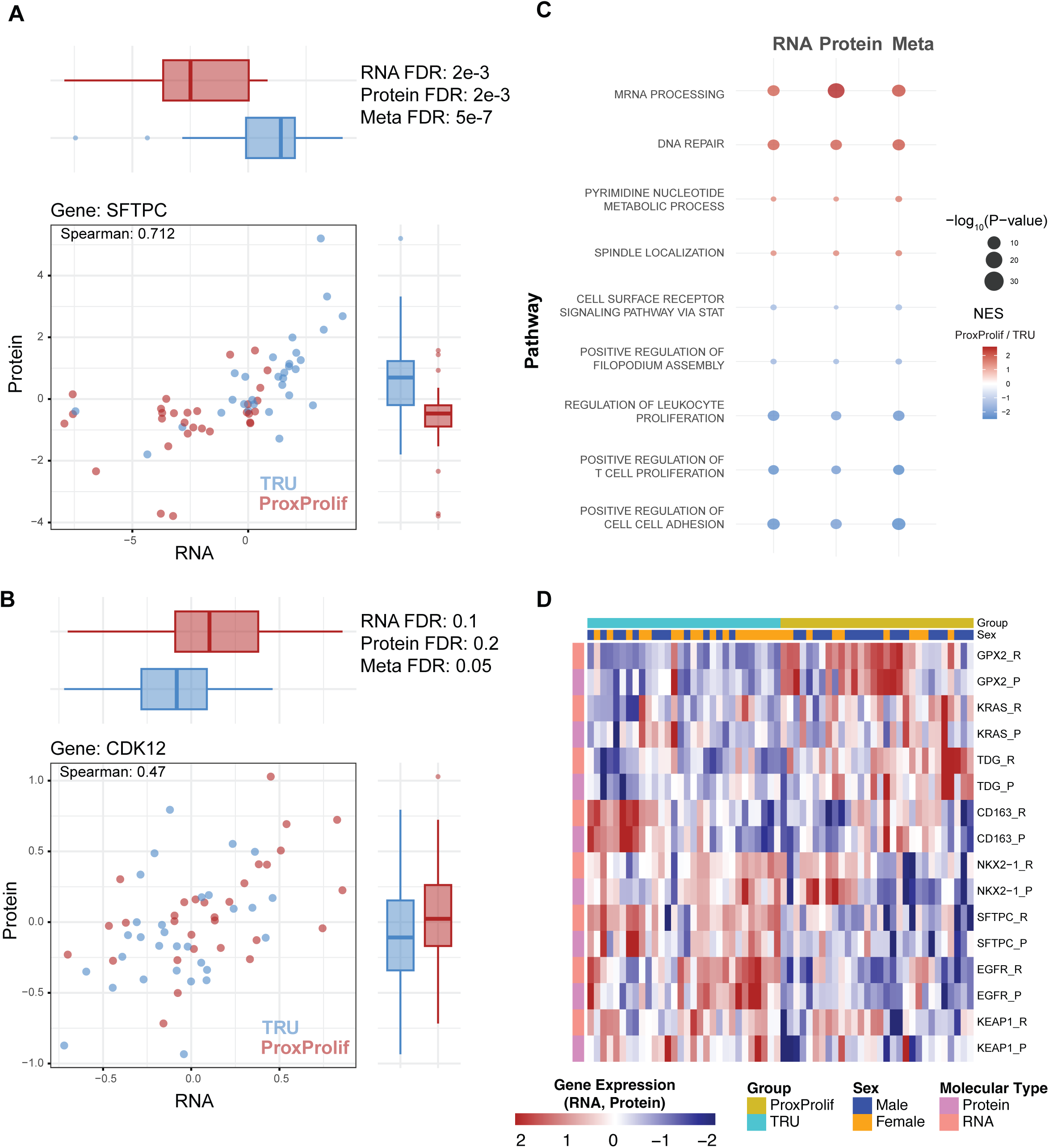
Data visualizations of rpGenePlot, rpPathway and rpHeatmap. Data visualizations for subtype differential expression in APOLLO cohort, related to Figure 2. For select genes, RNA and protein expression are displayed in a scatter plot with adjacent boxplots, to emphasize both the bivariate RNA and protein expression distribution and sample group collapsed differences. The Spearman correlation of RNA and protein also displayed (A, B). rpPathway displayed normalized enrichment scores (NES) and log p-values for the three models as a bubble plot, select pathways displayed, meta fdr < 0.05 (C). rpHeatmap displays interleaved RNA and protein expression for select genes of interest, fdr < 0.05 (D).

To enable functional interpretation of multi-omic DEGs, the rpPathway utility conducts gene set enrichment analysis on the RNA model DEGs, protein model DEGs, and meta model DEGs by calling FGSEA (43), resulting in overexpressed and underexpressed pathways associated with the sample groups or a continuous sample variable. rpMeta then creates an integrated bubble plot (44) to illustrate the pathway enrichments and log p-values by the three models (Figure 3C, Table S1C). In this way, rpPathway reveals pathways with stronger signal from the meta model compared to the RNA or protein models and vice versa.

To visualize the multi-omic expression of specific gene sets, the rpHeatmap utility generates an inter-leaved heatmap of RNA expression and protein expression to the same matched gene set (Figure 3D). Specifically, RNA and protein expression are normalized and organized as adjacent rows for the same genes and labelled by their molecular types, RNA or protein. rpHeatmap then calls pheatmap (45) to create this customized plot. The outcome is that patterns of RNA and protein expression can be easily visualized as pairs across key gene sets of interest. In this way, rpHeatmap effectively visualizes genes with highly correlated RNA and protein expression as well as genes with expression with greater contrast by RNA than by protein or vice versa. Similar heatmap visualizations are found in multi-omic studies such as CPTAC (46) and APOLLO (13); however, these publications utilized *ad hoc* code that is not available in an analysis package and is often regenerated per publication. rpMeta fills this gap for the analyst providing a complex visualization capability that is accessible through one function call.

### Benchmark datasets

Benchmark analyses were conducted using two published datasets of lung adenocarcinoma from CPTAC and APOLLO. APOLLO RNA-seq, MS-based proteomic expression data were retrieved from NCI’s Genomic Data Commons (GDC) portal (https://gdc.cancer.gov/about-data/publications/APOLLO-LUAD-2022) and Soltis et al (13) supplemental tables. CPTAC RNA-seq and proteomics data were downloaded from Gillette et al (17) supplemental tables. Expression subtypes for the CPTAC samples as defined in Soltis et al supplemental data were used for analysis. For benchmark analyses (Figure S1), we applied rpMeta to the APOLLO and CPTAC cohorts and used the limma method for both RNA and protein. We restricted analysis to genes commonly measured by RNA-seq and proteomics, with a maximum missing fraction of 50% (APOLLO n=7,467; CPTAC n=9,647; APOLLO and CPTAC n=6,839 genes) For the simulation and cross-cohort benchmarks, we restricted analysis to genes commonly measured by APOLLO and CPTAC cohorts’ RNA and protein, n = 6,839 genes.

## RESULTS

### rpMeta boosts sensitivity in simulation benchmark

To establish a simulated truth DEG set for benchmarking rpMeta, we leveraged the CPTAC cohort of RNA and protein expression to compare subtypes of lung adenocarcinoma, specifically, the Terminal Respiratory Unit (TRU) subtype and the Proximal-Proliferative (PP) subtype (Figure S1A) (17). We then calculated subtype DEGs in the CPTAC cohort by rpMeta using the limma model for both RNA and protein and selecting those genes with fdr < 0.001 (Table S1A). We defined the union of these RNA DEGs and protein DEGs as positives in our truth set. We then defined all other genes as negatives in the truth DEG set. Then to make a simulated test dataset, we utilized the APOLLO cohort of lung adenocarcinoma RNA expression, protein expression and subtype sample assignments (15). To create the simulated DEGs in APOLLO, we transformed the APOLLO expression data. Negative DEGs from the truth set were each randomly shuffled within gene in the simulated APOLLO cohort, thus removing the association of these genes with the subtype group of each sample and maintaining real-world expression distributions. Positive DEGs from the truth set were left as-is in both RNA and protein expression data in the APOLLO cohort. In this way, the simulated APOLLO data would preserve expression signal from RNA and protein for the positive genes, and the negative genes would be a random re-sampling of expression.

Then, we applied rpMeta to the simulated APOLLO data and compared performance of the RNA, protein, and meta model p-values to the truth DEG set using receiver operating characteristic (ROC) curve analysis (Figure 4A). The meta model displayed the largest performance with an area under the curve (AUC) of 0.912 compared to the RNA AUC of 0.851 and the protein AUC of 0.827. The ROC curve analysis also revealed more true positive DEGs by the meta model than the RNA and protein models across the range of false positive DEGs. Therefore in simulated real-world multi-omic data, the meta model utilizing the joint signal of RNA and protein had greater sensitivity.

**Figure 4:**
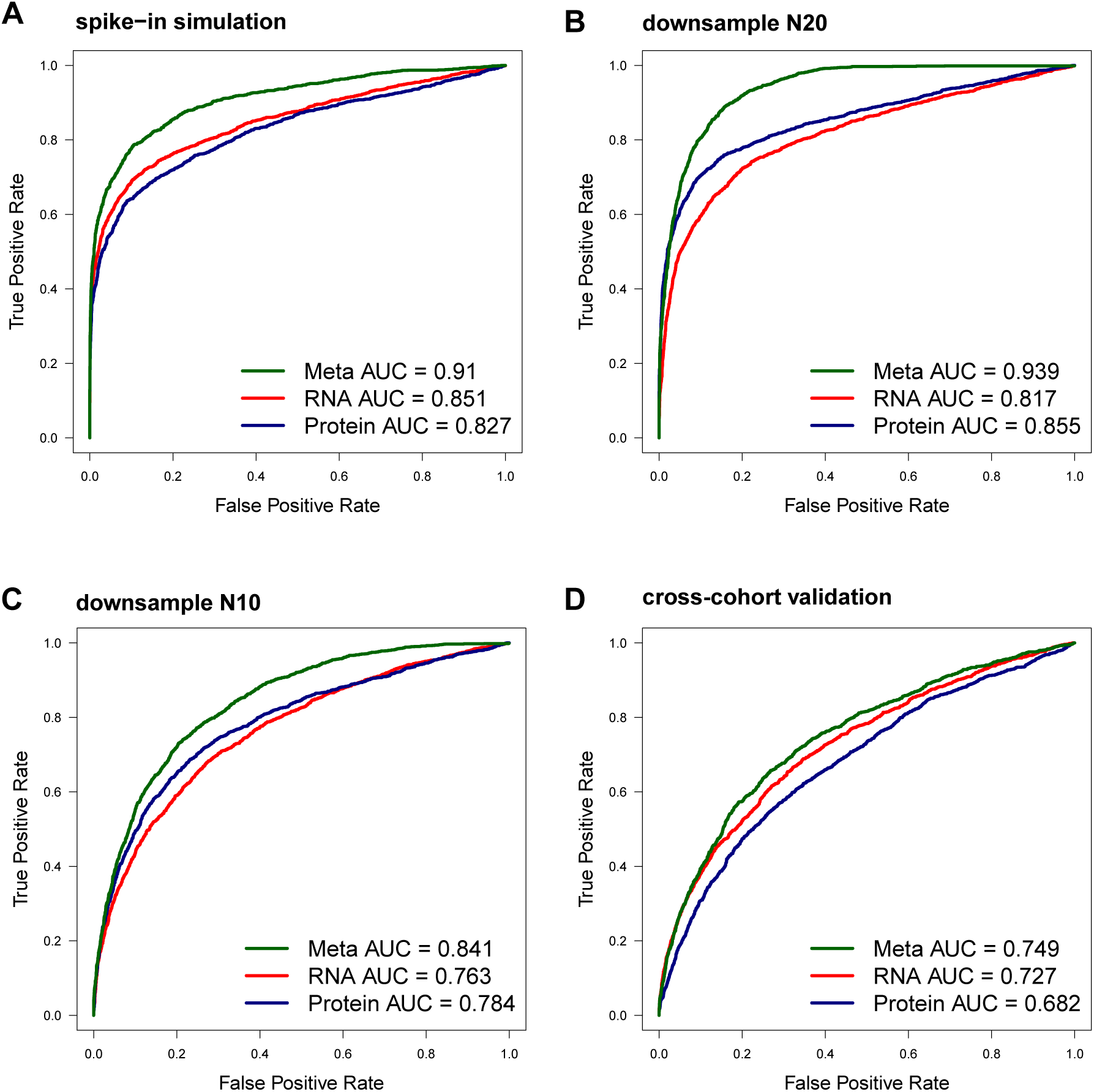
rpMeta benchmark analyses. Receiver operating characteristic curves and area under the curve (AUC) are presented for four benchmark analyses (Figure S1). Line color indicates the DEG model from rpMeta, following to the legend.

### rpMeta displays greater sensitivity in downsampling benchmark

To determine if RNA and protein integration for DEG analysis was beneficial in the small sample size setting, we performed a downsampling benchmark (Figure S1B). First, we defined a truth DEG set using the full APOLLO cohort of lung adenocarcinomas containing 30 TRU subtype samples and 30 PP samples, restricted to genes commonly measured by RNA and protein (Table S1B). rpMeta was run to detect RNA and protein DEGs between TRU and PP. Positive DEGs in the truth set were genes with a fdr threshold of 0.01 in either RNA or protein. Negative DEGs in the truth set were all other genes. The first subsample test set (N20) was a randomly selected subset of 20 TRU samples and 20 PP samples from the full APOLLO cohort. rpMeta was run on N20 test set to detect RNA, protein and meta DEGs. In the N20 test set, the meta model had the greatest performance with an AUC of 0.939 compared to an RNA AUC of 0.817 and a protein AUC of 0.855 (Figure 4B). We then created another test set (N10) by subsampling 10 TRU samples and 10 PP samples from the full APOLLO set. rpMeta was run on the N10 test set with the same options as the N20 set. In the N10 test set, all models had reduced performance due to the smaller sample size; however, the meta model continued to display the greatest performance compared to RNA and protein models with AUCs of 0.841, 0.763, and 0.784 respectively (Figure 4C). Similar to the spike-in benchmark, the ROC curves showed more true positive DEGs by the meta model than the RNA and protein models across the range of false positive DEGs. Therefore through downsampling benchmark analysis, the meta model of rpMeta had greater sensitivity compared to RNA and protein models.

### rpMeta yields greater performance in cross-cohort validation

To benchmark the performance of rpMeta between two real world datasets, we utilized the full datasets of CPTAC and APOLLO samples using all PP and TRU samples as a cross-cohort benchmark (Figure S1C). First in the CPTAC cohort, we calculated DEGs using rpMeta. Positive genes for the truth set were defined as those with a fdr < 0.05 by the RNA or protein model (Table S1A). Negative genes for the truth set were all other genes. Then, we used all APOLLO TRU and PP samples as a test set. DEGs by the RNA, protein and meta models were calculated using rpMeta in the test set (Table S1B). Comparing the APOLLO DEGs to the CPTAC truth DEG set, we found that the meta model had the greatest performance with an AUC of 0.749 compared to smaller AUCs in the RNA and protein models of 0.727, and 0.682 respectively (Figure 4D). In summary, the rpMeta model outperformed the RNA and protein models in this cross-cohort validation.

### rpMeta discovers differentially expressed genes and pathways in lung cancer

To demonstrate how rpMeta may advance biological interpretation of a real-world multi-omics use case, we applied rpMeta to the full APOLLO cohort of 30 TRU and 30 PP with a fdr threshold of 0.05. The RNA and protein models detected similar numbers of DEGs (n = 2,193, n = 2,081, respectively) (Figure 2A, Table S1B). The meta model detected a larger number of DEGs (n = 3,385). Many (31%) of the meta DEGs were also detected by both the RNA and protein models. For example, surfactant protein C, *SFTPC*, was overexpressed in the TRU subtype by RNA and protein, consistent with being a known strong marker of this subtype from earlier studies (30) (Figure 3A). Another 31% of meta DEGs were detected by the RNA model and not the protein model. Another 27% of rpMeta DEGs were detected by rpMeta protein and not by rpMeta RNA.

Strikingly, a much larger number of DEGs, n = 376, were detected only by meta model compared to DEGs only detected by RNA (n = 110) or protein (n = 110) models. These 376 novel DEGs represent genes that would not be detected by analyzing RNA-seq or MS-based proteomics alone. Review of the DEGs unique to the meta model revealed many were relevant to the biology of lung adenocarcinoma, including cyclin-dependent kinase 12, *CDK12*. *CDK12* was only significantly detected by the meta model and was overexpressed in the PP subtype, as displayed in the rpGenePlot (Figure 3B). *CDK12* is involved with cell cycle machinery, required for proliferation in lung cancer, and is involved in DNA repair (47), all of which are consistent with the PP subtype’s known characteristics (13,48).

Beyond genes, the meta model revealed new biological processes in the subtypes (fdr < 0.05) not detected by the RNA or protein models (fdr > 0.05) (Table S1C). For example, the spindle localization and pyrimidine nucleotide metabolic processes were overexpressed in the PP subtype. These are generally consistent with the known overall high cellular proliferation pathway activity of the PP subtype (48); however, these novel pathway enrichments detected only by the meta model reveal detailed biological processes for deeper characterization of the PP subtype and serve as leads for future experimentation. Similarly for the TRU subtype, the meta model detected pathways not found by the protein or RNA models. Specifically, the TRU subtype overexpressed biological processes of positive regulation of filopodium assembly and cell surface receptor signaling via STAT (Figure 3C). This is consistent with the TRU subtype’s known high expression of lung development processes and immune cell presence (13); however, the meta model has now identified new specific biological processes that characterize TRU more deeply. This real-world use case demonstrates that rpMeta discovers new knowledge in the form of genes and pathways compared to RNA and protein models, thus advancing biological discovery.

To visualize aggregate DEGs from these analyses, the rpHeatmap displays user-specified DEGs overexpressed in each subtype by RNA and by protein at sample-level resolution (Figure 3D). rpHeatmap enables readers to see genes that have high consistency between RNA and protein and differential expression such as *SFTPC* in the TRU subtype and *GPX2* in the PP subtype, emphasizing the robustness of these genes as markers for the subtype across both RNA and protein. rpHeatmap also reveals genes and samples in which RNA or protein is leading in relative differential expression, even within the group of interest, such as in *EGFR* and in *KRAS* in which a few samples have greater expression by protein than RNA. These individual samples could be related to unique features, such as somatic mutations, which the user could also display via rpHeatmap.

## DISCUSSION

To determine if combining RNA-seq and MS-based proteomics for DEG analysis increased performance, we developed rpMeta as a novel method and R package. For the first time, we demonstrated that combining RNA-seq and MS-based proteomics increases sensitivity in detecting DEGs across the large co-measured gene space of RNA-seq and MS-based proteomics through three benchmark analyses. Our innovative spike-in simulation benchmark demonstrated that the meta model achieves superior sensitivity to both the RNA and protein models. Our downsampling benchmark demonstrated that rpMeta has a large performance gain in small sample sets. rpMeta may be helpful for future investigators who may have small cohorts and who are seeking to boost their power to detect DEGs. These results may also be helpful for investigators seeking to balance their experimental designs and costs between adding more samples or adding a second ‘omic platform, RNA-seq or proteomics. Finally, we demonstrated that rpMeta has the greatest sensitivity in detecting real world DEGs by holding one lung cancer cohort, CPTAC, as truth and another lung cancer cohort, APOLLO, as the test set.

To demonstrate in practice if rpMeta would enable new DEG discoveries compared to RNA or protein alone, we applied rpMeta to the entire APOLLO cohort comparing the PP and TRU subtypes as a real-world use case. The meta model detected more DEGs and pathways than the RNA or protein models providing more signal for biological characterization. We described that these genes, e.g. *CDK12*, and pathways, e.g. spindle localization, are consistent with and expand the overall molecular characteristics of the subtypes from prior literature. All together, these real-world results demonstrate that the gain in joint RNA-seq and proteomics DEG analysis not only increases statistical significance but also increases biological discovery.

In addition to the DEG calculations, rpMeta provides four data visualization utilities for multi-omic DEG analysis. These novel visualizations enable the analyst to interpret and explore the integrated expression data in detail and produce publication quality figures for the manuscript reader to interpret. rpMeta follows best practices for bioinformatics software development (49–51). We utilized standard data formats in gene expression matrices and sample information table, and write all options to log text files. rpMeta provides a single driver function that wraps all analyses together. rpMeta also includes utility functions for direct calculation of specific components of the workflow. To maximize accessibility and transparency, we wrote rpMeta as an R package and made it available open source on github (https://github.com/zhangx8/rpMeta). rpMeta includes technical documentation and a detailed tutorial so that rpMeta will be accessible to a range of analysts from students through expert power users. Future development of rpMeta may focus on expanding rpMeta to additional types of analysis beyond the two group and continuous variable comparisons, the latter of which we utilized in a large pan-cancer study (52). Existing multi-omic tools, such as mixOmics (53) and linkedOmics (54), focus on multi-omic data integration for analytical purposes other than differential expression. rpMeta is the first multi-omic tool specialized for multi-omic DEG analysis by meta-analysis, and the first package including multi-omic DEG visualizations, such as the integrated rpHeatmap. Our report here is the first performance evaluation of DEG meta-analysis of RNA-seq and MS-based proteomics. Therefore, rpMeta uniquely fills a current gap not possible with existing tools.

In summary, we demonstrate, for the first time, that integrating RNA-seq and proteomics boosts sensitivity to detect DEGs compared to RNA or protein alone. We developed rpMeta as an open-source tool, filling a gap in the bioinformatics community to address a new analysis task – meta-analysis of RNA and proteomics for DEG analysis. rpMeta with its novel multi-omic data visualization and detailed tutorials is accessible to a wide range of analysts and conducts multi-omic DEG analysis in a standardized way which will lead to increased analysis reproducibility and time savings for the analyst with publication-quality figures and result data tables. Through its increased sensitivity, rpMeta derives new power from multi-omics data for DEG analysis, advancing biological discovery.

## DATA AVAILABILITY

This study analyzes publicly available datasets from APOLLO and CPTAC cohorts available at NCI’s Genomic Data Commons and Proteomic Data Commons. The utilized versions of these data are provided in the data of the R Package at https://github.com/zhangx8/rpMeta).

## SUPPLEMENTARY DATA

Supplementary Data are enclosed with the manuscript.

## ACKNOWLEDGEMENTS

We thank the testers of rpMeta software. The authors acknowledge contributions of anonymous patients to the previously published cohorts utilized in this study. The opinions and assertions expressed herein are those of the author(s) and do not reflect the official policy or position of the Uniformed Services University of the Health Sciences or the Department of War. References to non-Federal entities or products do not constitute or imply a Department of War or Uniformed Services University of the Health Sciences endorsement.

## AUTHOR CONTRIBUTIONS

MDW: conceptualization, formal analysis, writing – original draft, visualization, funding acquisition, software, methodology, investigation. XZ – formal analysis, software, writing – original draft, investigation, visualization, data curation; JL: investigation, formal analysis, writing – review & editing; RLK: supervision, writing – review & editing, CLD: Funding acquisition, investigation, resources, writing – review & editing. CDS – resources, writing-review & editing, supervision

## FUNDING

Funding support was provided in part from USUHS award #HU0001230028 to the Murtha Cancer Center Research Program and USUHS award #HU00012420045 to the Center for Military Precision Health.

Supplementary Material for Zhang X. et al (2026) **Combining RNA-seq and proteomics boosts sensitivity to detect differentially expressed genes by rpMeta**

---

**Figure S1:**
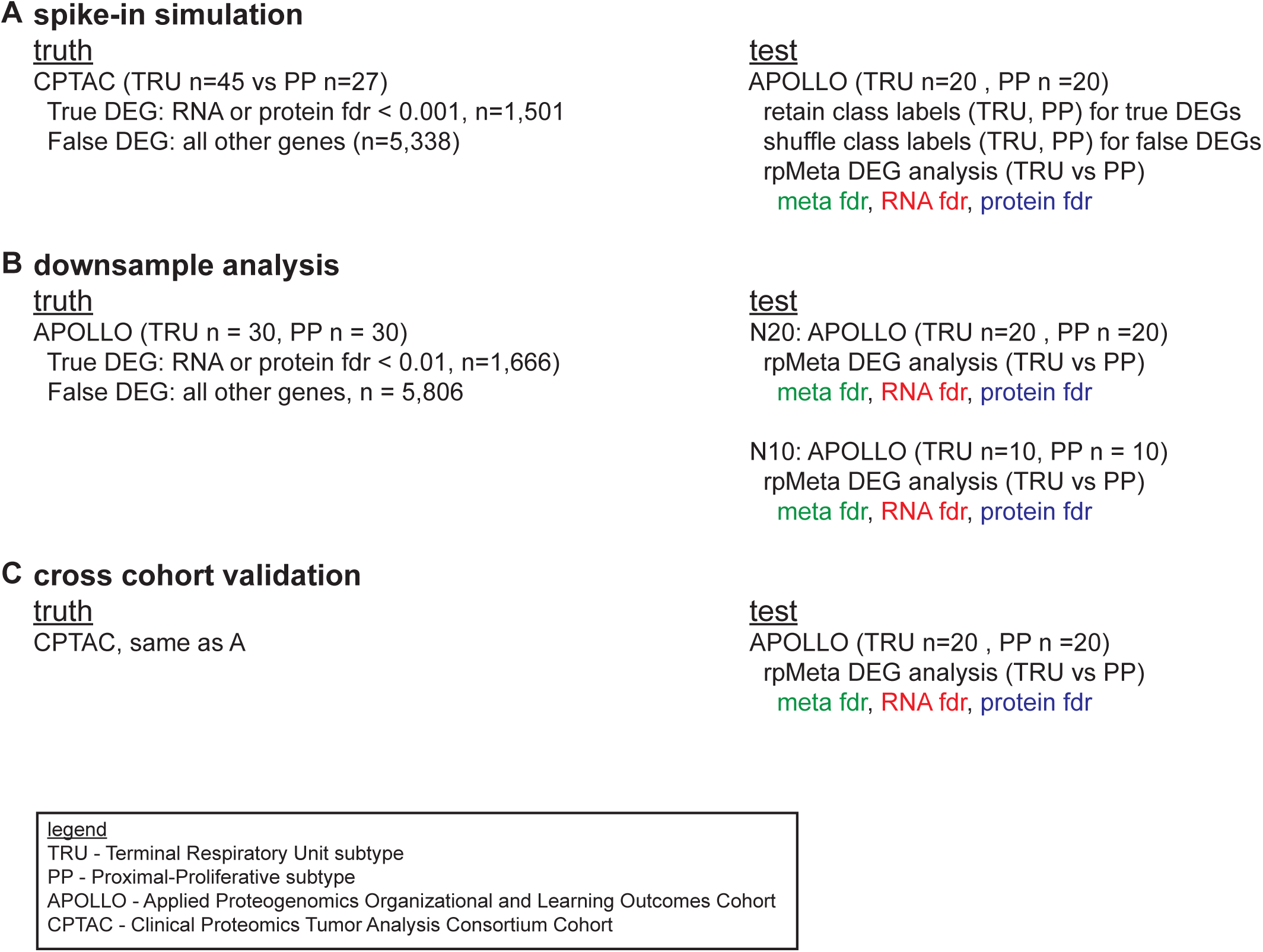
Benchmark analysis design for rpMeta. The datasets, samples sizes and options utilized for truth and test sets are presented for three benchmark analyses: spike-in simulation, downsample analysis, and cross cohort analysis. fdr color code refers to Figure 4.

**Table S1:** rpMeta results. Differentially expressed genes between TRU and PP subtypes in full CPTAC cohort (A) and full APOLLO cohort (B). Fold change > 0 indicates overexpression in PP subtype vs TRU subtype. rpPathway results for enriched gene ontology biological processes in the APOLLO cohort (C). Enrichment score > 0 indicates overexpression in PP subtype vs TRU subtype. Provided as a separate file.

